# Pancreatic cancer metastasis is regulated by an eleven amino-acid sequence

**DOI:** 10.64898/2025.12.07.692837

**Authors:** Guillem Fuertes, Daniele Di Biagio, Riddhi P Sontakke, Eleni Maniati, Benjamin Jenkins, Gareth J Thomas, John F Marshall

## Abstract

Pancreatic ductal adenocarcinoma (PDAC) has a very poor prognosis with a 5-year survival rate less than 5% because of its ability to metastasise, its late detection and the lack of effective therapies. The Integrin αvβ6 is highly overexpressed in PDAC and correlates with poor prognosis. The integrin β6 subunit contains a unique C-terminal tail of 11 amino acids (aa) that regulates downstream signals, although the mechanism remains unclear. Here, using integrin β6-deficient cells lines, we have developed two PDAC mouse models overexpressing the full-length β6 and a mutant β6 lacking the C-terminal 11aa (αvΔβ6). In vitro, αvβ6 overexpression increased cell proliferation, migration and invasion in a 3D spheroid model. The elimination of the C-terminal 11aa decreased proliferation and totally impaired cell migration and invasion. αvΔβ6 cells also expressed reduced MMP’s in vitro. In vivo, orthotopic implantation of αvβ6 overexpressing cells showed decreased overall survival and more spontaneous metastasis compared to αvΔβ6 and αvβ6-null cells. Therefore, the C-terminal 11aa of the integrin β6 subunit regulates PDAC progression and metastasis.

## Introduction

The 5-year survival from pancreatic ductal adenocarcinoma (PDAC) remains at <5% [1] so identifying novel effective therapeutics is critical. The major issue with PDAC is late presentation due to limited clinical signs resulting in a large fraction of patients being diagnosed with late-stage disease including distant spread, and it these metastases that eventually lead to the death of the patients [2]. Thus, until we have developed effective methods to detect PDAC earlier, we must treat or prevent formation of PDAC metastases if we are to improve overall survival.

The integrin αvβ6 is not usually expressed at detectable levels in most tissues but it becomes upregulated in many cancers including colon, stomach, lung, breast, ovary, cervix, skin, where its expression correlates with poor survival [3-9]. Our group and others have reported that the integrin αvβ6 is overexpressed in most (>90%) human PDAC tumours compared with the healthy pancreas which had little or no detectable αvβ6. Importantly the increased αvβ6 expression was maintained in the matched metastases [10] and high levels of αvβ6 protein expression and β6 mRNA correlated with poorer survival from PDAC [10, 11]. Since αvβ6 is associated with more advanced disease and poorer survival in PDAC it may suggest that it is actively promoting metastasis.

In earlier studies we reported that αvβ6 induced oral carcinoma cell invasion, in part at least, by causing the increased secretion of the matrix metalloproteinase zymogens pro-MMP2 and pro-MMP9 [12]. Mechanistically we further showed that the C-terminal 11 amino acids on the cytoplasmic tail of β6, which is present only in the β6 integrin-subunit (Fig 1A), regulated the increased release of the pro-MMPs [13]. Studies by others reported the same 11 amino-acids correlated with increased cell proliferation [14-16] although again mechanisms were unclear.

**Figure 1.**
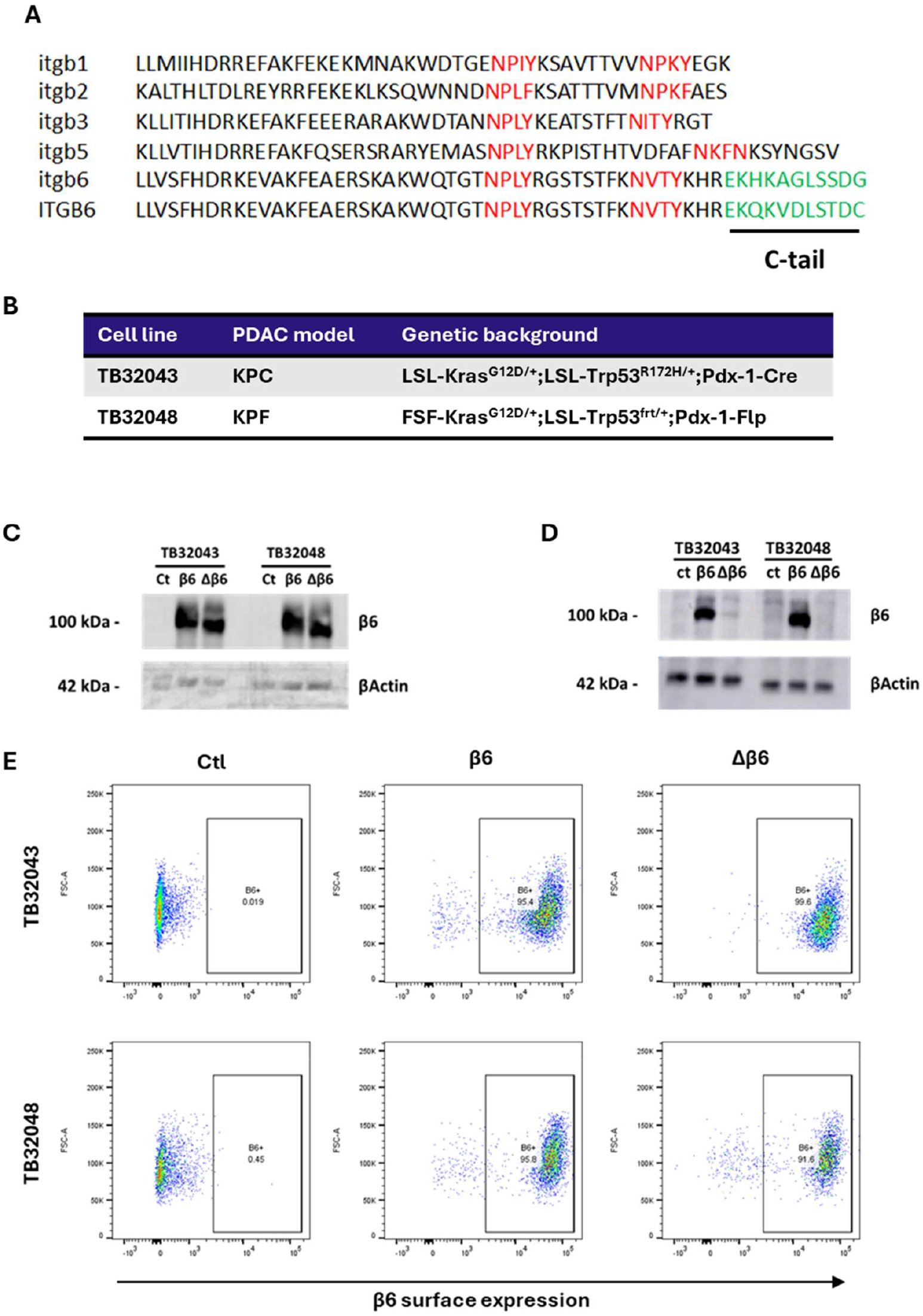
Development of a KPC- and KPF-derived PDAC models overexpressing integrin β6. (A) Comparison of human and mouse integrin (ITGB6 and Itgb6 respectively) cytoplasmic tail sequences, with other mouse integrin β subunits. Note the additional 11 amino-acid extension on the ITGB6 and Itgb6 (B) Table detailing the genetic background of the KPC- and KPF-derived PDAC cell lines used in this study. (C) Ctl, β6- and Δβ6-expressing TB32043 and TB32048 cells were analysed by western blotting with a goat anti-β6 antibody recognising the extracellular N-terminal region of the protein. (D) Ctl, β6- and Δβ6-expressing TB32043 and TB32048 cells were analysed by western blotting with a goat polyclonal anti-β6 antibody recognising the C-terminal end of β6. (E) Ctl, β6- and Δβ6-expressing TB32043 and TB32048 cells were analysed by flow cytometry to determine integrin localisation at the membrane. All cells were labelled with anti-αvβ6 antibody 10D5.

Here we describe that the integrin β6 cytoplasmic tail 11 amino-acids is key, not only for the integrin αvβ6-regulated increase in proliferation, migration and invasion in vitro, but is essential for lung metastasis of PDAC.

## Results

### 1. Development of integrin αvβ6-overexpressing PDAC murine models

To determine how αvβ6 drives PDAC progression and metastasis and study precisely what role the β6 C-terminal tail plays in αvβ6 dependent functions, we have generated a cellular model in which the full length wild-type murine β6 subunit or a mutant form lacking the last 11 aa (hereafter Δβ6) have been overexpressed in PDAC murine cell lines that lack the expression of endogenous β6.

Using lentiviral vectors, we transduced the wild-type or truncated murine β6 sequences (itgb6) (or the empty vector) into cell lines derived from two genetically different PDAC models: the KPC-derived cells TB32043 [17] and the KPF-derived cells TB32048 [18] (Fig 1B). Using fluorescence activated cell sorting we selected two different isogenic panels of three cell lines that expressed no αvβ6, wild-type αvβ6 or the αvΔβ6.

Similar levels of expression of β6 and Δβ6 in TB32043 and TB32048 cell models were confirmed by western blotting (Fig 1C). Additionally, the expression of Δβ6 was verified by western blotting using an anti-integrin β6 goat antibody that binds to the C-terminal end of the protein (Fig 1D), showing no protein detection on the Δβ6 cells samples. The surface expression of both, β6 and Δβ6, was measured by flow cytometry using an N-terminal recognising integrin β6 antibody. Data confirmed that more than 90% of cells show similar levels of β6 and Δβ6 in the cell membrane (Fig 1E).

### 2. Expression of β6 induces increased proliferation and migration through its C-terminal 11 amino-acid tail

Expression of integrin αvβ6 has been described to promote proliferation and migration in various cancers [10, 19]. The role of the β6 C-tail in controlling proliferation was firstly described in colon carcinoma cells in 1996 [15]. Therefore, we investigated how the deletion of the 11 aa from the β6 C-tail affects proliferation in murine PDAC cell lines.

β6 overexpression in TB32043 and TB32048 cells significantly (p>0.001) increased cell proliferation after 72h (Fig 2A). Deletion of the β6 C-tail (Δβ6) significantly (p<0.05) decreased cell proliferation compared with wild-type β6-expressing cells (Fig 2A). Consistently, when cells were treated with integrin αvβ6-blocking antibodies 264RAD and 10D5, cell proliferation significantly decreased (p>0.01-0.001) in all β6-expressing cells, while only a small decrease was observed in Δβ6-expresing cells, reaching significance only in the TB32048 Δβ6 cells (Fig 2B). Treatment with αvβ6-blocking antibodies did not affect control (Ctl) cells proliferation (Fig 2B).

**Figure 2.**
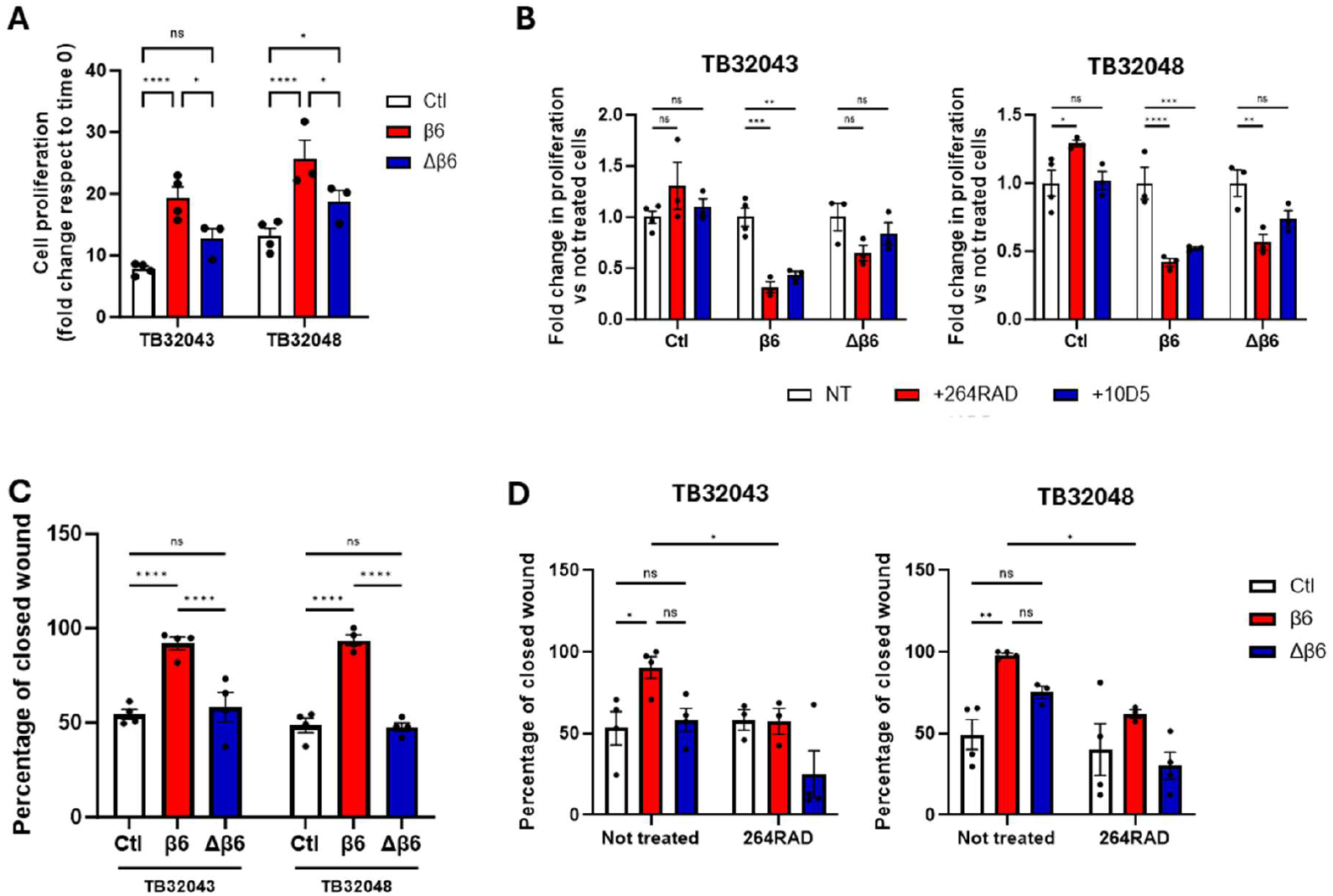
Integrin αvβ6 overexpression increases cell proliferation and migration through its cytoplasmic tail. (A-B) Cell proliferation of TB32043 and TB32048 cell lines was measured in Incucyte for 72h. Cells were cultured in DMEM 10% FBS. (A) Proliferation was quantified using cell confluency and fold change respect to time 0 (Mean ± SD of four biological replicates). (B) Cells were treated with 10D5 or 264RAD as indicated. Proliferation was quantified using cell confluency at 72h and fold change respect to untreated conditions (Mean ± SD of three biological replicates). (C-D) TB32043 and TB32048 cell migration was measured in a scratch-wound healing assay. Migration is shown as percentage of the closed wound. In (D) cells were treated with 264RAD as indicated (Mean ± SD of at least three biological replicates).

Cell migration was also evaluated using scratch-wound healing assays in TB32043 and TB32048 cells. Cells expressing β6 showed a significant increase in migration, which was impaired significantly (p>0.001) in Δβ6-expressing cells (Fig 2C). 264RAD treatment significantly decreased cell migration in TB32043 and TB32048 β6 cells, while no significant effect o fth eantibody was seen in Ctl or Δβ6 cells (Fig 2D).

### 3. Integrin β6 cytoplasmic tail promotes PDAC cell invasion

The integrin β6 is implicated in controlling cell invasion [10, 13]. We measured tumour cell invasion of the TB32043 and TB32048 panels of cell lines using a 3D spheroid model [20]. Spheroid invasion was tested in spheroids containing only PDAC cancer cells (Fig 3A) or in combination with pancreatic stellate cells (PSCs) (Fig 3D). In the mono-culture spheroids, expression of β6 significantly (p<0.001) increased the spheroid invasive area, which was reduced to background control (Ctl) levels when the β6 C-tail was deleted in Δβ6-expressing cells (Fig 3B-C).

**Figure 3.**
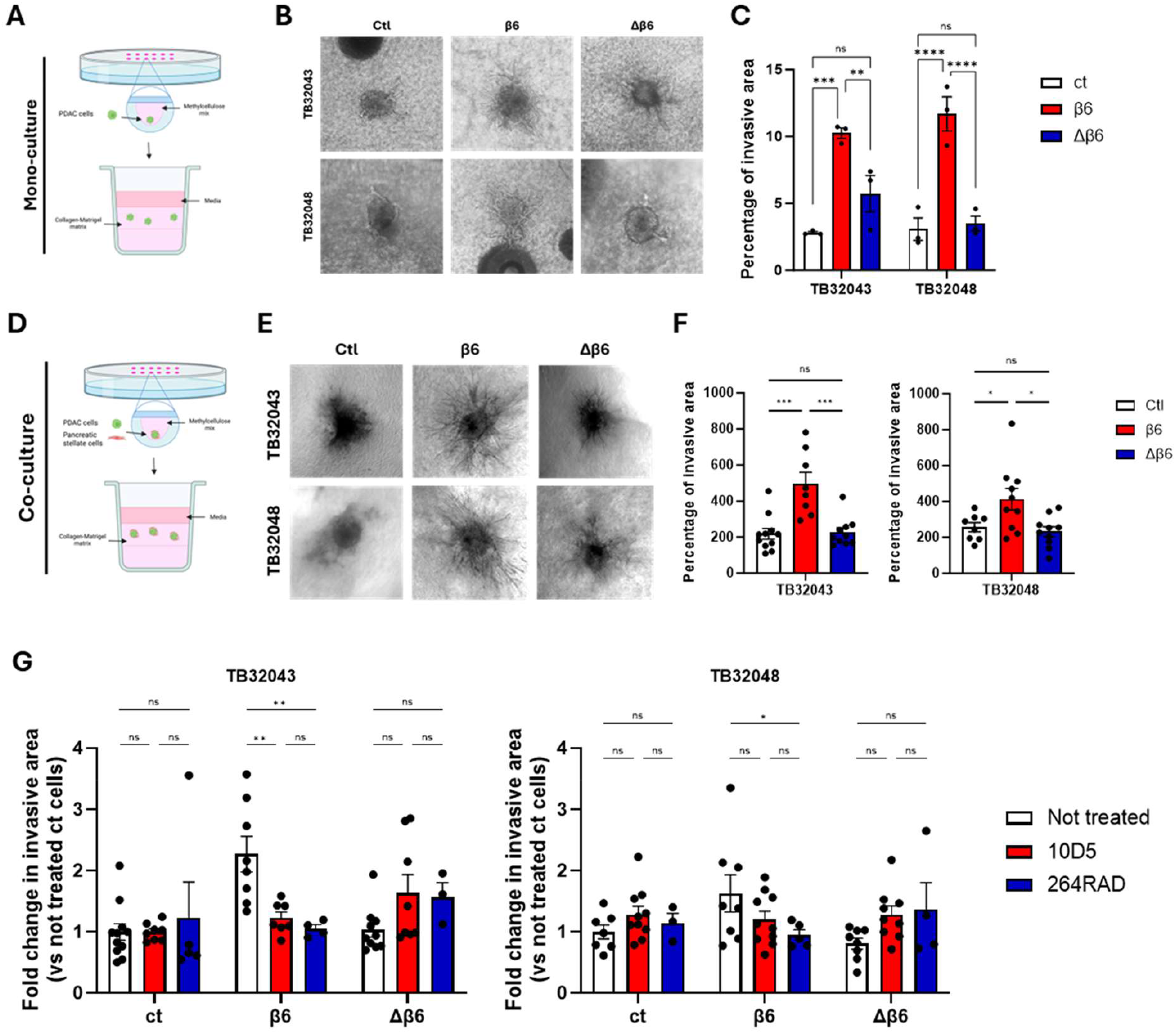
Integrin αvβ6-induced PDAC invasion is impaired after cytoplasmic-tail 11 amino-acid deletion. (A) Schematic view of mono-culture 3D spheroids invasion assay. (B) Representative images of Ctl, β6- and Δβ6-expressing TB32043 and TB32048 spheroids. (C) Quantification of spheroids invasive area (Mean ± SD of three biological replicates). (D) Schematic view of co-culture (PDAC cells + PSCs) 3D spheroids invasion assay. (E) Representative images of Ctl, β6- and Δβ6-expressing TB32043 and TB32048 cells co-cultured with PSCs in spheroids. (F) Quantification of co-culture spheroids invasive area (Mean ± SD. Dots represent individual spheroids from three biological replicates). (G) Quantification of co-culture spheroids invasive area treated with 10D5 and 264RAD. Invasive area is represented as fold change vs untreated Ctl cells (Mean ± SD. Dots represent individual spheroids from three biological replicates).

Similar data were generated with the co-culture spheroids. PDAC cells were combined with PSCs in a 1:2 ratio to generate spheres (Fig 4D). In the co-culture experiments, the expression of β6 also resulted in a significant (p<0.05) increase in invasion (Fig 3E-F) whereas spheroids containing Δβ6-expressing cells showed significantly reduced invasive capacities, migrating similarly to Ctl cells spheroids (Fig 3E-F).

**Figure 4.**
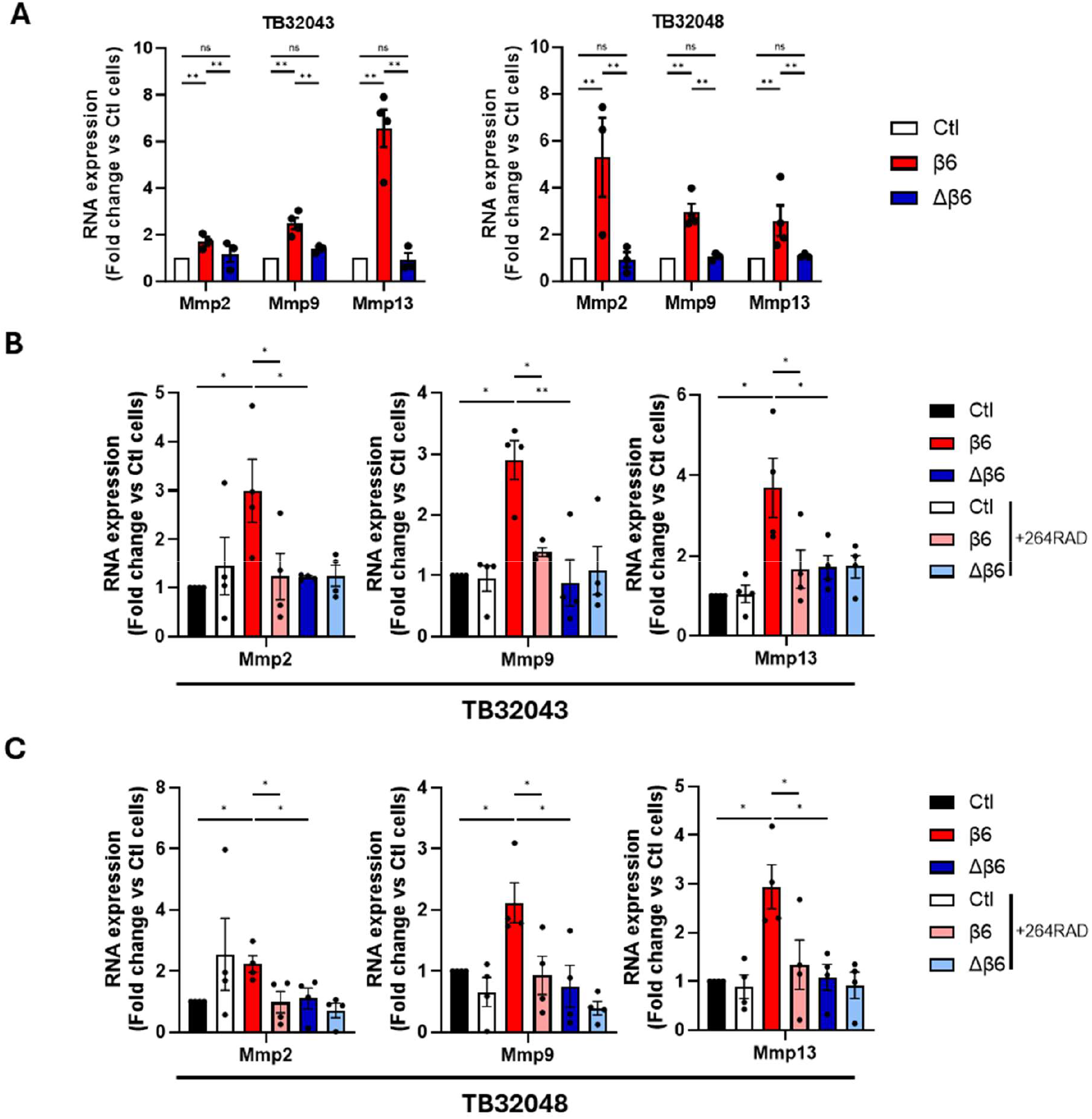
Integrin β6 C-tail 11 amino-acids regulates Mmp2, Mmp9 and Mmp13 gene expression. RNA was isolated from TB32043 and TB32048 Ctl, β6- and Δβ6-expressing cells (A) or cells treated with 264RAD (B-C) and expression of Mmp2, Mmp9 and Mmp13 was analysed by qRT-PCR. Data are represented as fold change relative to Ctl cells levels for each gene (Mean ± SD of at least three biological replicates performed in triplicates).

Spheroid treatment with αvβ6-blocking antibodies 264RAD or 10D5 both significantly (p<0.01) reduced spheroid invasion in β6-expressing cells, while no effect was observed in the Ctl and Δβ6-expressing cells (Fig 3G). Together, these results suggest that the β6 C-terminal cytoplasmic 11 amino-acids has a key role in regulating β6-dependent 3D invasion.

### 4. The β6 cytoplasmic tail regulates expression of matrix metalloproteinases genes

Previous studies have shown that the integrin β6 cytoplasmic tail promotes tumour cell invasion in human oral squamous cell carcinoma by controlling secretion of matrix metalloproteinases [13]. Therefore, the dependency of MMP expression on the β6 cytoplasmic tail in PDAC was investigated.

Using qRTPCR, RNA expression of Mmp2, Mmp9 and Mmp13 was significantly increased in wild-type β6-expressing cells compared to Ctl cells (Fig 4A). Consistent with previous observations, expression of Mmp2, Mmp9 but also Mmp13 was affected by β6 cytoplasmic tail deletion of the C-terminal 11 amino-acids, as observed in Δβ6-expressing cells (Fig 4A). Similarly, when β6-expressing cells were treated with 264RAD, the RNA levels of Mmp2, Mmp9 and Mmp13 were significantly decreased (Fig 4B-C) suggesting that MMP gene expression may depend on ligand-binding to αvβ6.

### 5. The β6 C-terminal 11 amino-acids regulates lung metastasis and survival

We transduced the TB32043 and TB32048 panels of cell lines with luciferase (Addgene) and then injected 1000 cells from each of lines orthotopically into the pancreas of C57Bl/6 mice (Fig 5A). After 7 days tumours were detectable. Figure 5B and C shows that expression of wild-type β6 significantly reduced survival (p<0.001) compared to the β6-deficient Ctl cells. Interestingly the Δβ6-expressing cells exhibited an intermediate survival response (Fig 5B-C).

**Figure 5.**
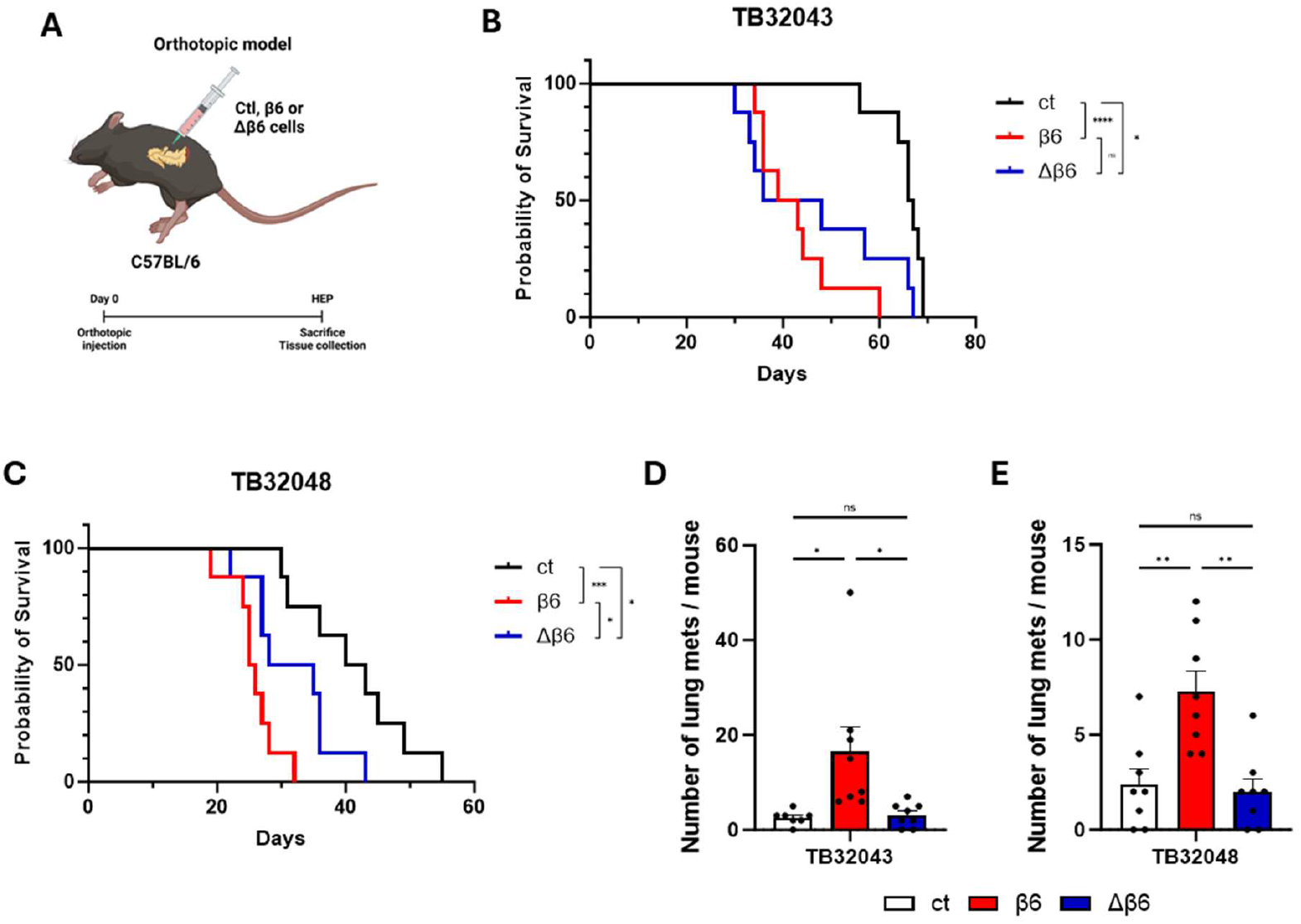
Integrin β6 C-tail 11 amino-acids reduce survival and promote metastasis. (A) Schematic view of the orthotopic PDAC model. After mice termination, tissues were collected, fixed and stained for H&E. (B-C) Kaplan Meier curves show β6 expression reduces survival compared with Ctl and Δβ6-expressing cells in TB32043 (B) and TB32048 (C) cell models (n=8 mice per condition). (D-E) Lungs were analysed after staining with H&E for metastases. Data is represented as the number of lung metastases per mouse (n=8 mice per group).

When lungs from the mice were harvested it was clear that the β6 C-terminal 11 amino-acids was essential for the αvβ6-dependent increase in metastasis (Fig 5D-E). Thus, lung metastasis was significantly (p<0.05) higher in the β6-expressing TB32403 and TB32048 cell lines compared with either the β6 deficient Ctl cells or the Δβ6-expressing cells.

## Discussion

The expression of integrin αvβ6 is increased in most human PDAC tumours and correlates with poor overall survival [10, 11]. This increased expression makes the integrin αvβ6 a promising candidate for targeted therapies as the normal pancreas expresses very little αvβ6 [21]. The implications of αvβ6 overexpression have been widely studied by us and others, showing that integrin αvβ6 expression promotes higher proliferation and invasion in different types of cancer cells [8, 19, 22, 23].

In this study we have also demonstrated that expression of β6 in murine PDAC cells promotes an increase in cell proliferation, migration and invasion (Figures 2 and 3). These results match with previous observations in other type of cancers [24-27]. In particular, here we describe that the β6 cytoplasmic tail is required for β6-dependent cell proliferation, migration and invasion of PDAC cell lines. We have shown that PDAC cells expressing β6 that lacks its cytoplasmic 11aa-tail no longer exhibit increased proliferation, migration and invasion. The β6 C-tail deletion not only abolished PDAC mono-culture spheroids invasion, but also in spheroids where PDAC cells were co-cultured with pancreatic stellate cells, suggesting that the β6 C-tail may also regulate the crosstalk between cancer cells and stellate cells.

However, although us and others have demonstrated that β6 C-tail controls β6-dependent functions, the mechanism of action still remains not clear. One possibility is that β6 C-tail deletion could affect its localisation at the cell membrane, but as shown in our study, Δβ6 localises at the membrane. Also, it has been observed that C-tail deletion doesn’t affect β6 localisation to focal adhesions [28]. Previous work in the lab has shown that β6 promotes increased production of matrix metalloproteases in human oral squamous cell carcinoma [13]. Here we also demonstrate that β6 C-tail induces the expression of several Mmp genes including Mmp2, Mmp9 and Mmp13, in PDAC. The increased Mmp gene expression may be implicated in promoting the observed increased invasion, although it may not be the only mechanism.

As invasion is an integral part of the metastatic process and as metastasis is the key reason for reduced survival from cancer, we also tested whether the C-terminal 11aa of β6 was involved in the metastatic capacity of PDAC cells and of survival. Figures 5B and C show that αvβ6 promotes significantly reduced survival in both TB32043 and TB32048 PDAC cell line models, which likely is due in part to the increase in lung metastasis (Figure 5D and E). The data also show that the C-terminal 11aa of β6 are essential for the increased lung metastasis and have an intermediate effect on survival in both PDAC models.

The mechanisms of how the C-terminal 11aa of β6 regulates metastasis are not clear. Certainly, the regulation of Mmp genes may partly explain these observations. One could speculate that the cytokine TGF-β may also be involved.

The cytokine TGF-β is implicated strongly in promoting the metastatic behaviour of cancer cells by a variety of mechanisms including inducing epithelial-to-mesenchymal transition [29]. As the integrin αvβ6 can activate latent-TGF-β [30] this could explain the observed effects on metastasis. The αvβ6 recognizes and binds to RGD motifs found in the latency associated peptide (LAP) [31] that surrounds and protects the TGF-β homo-dimer. αvβ6 activates TGF-β through a force-mediated mechanism, binding to LAP and transducing force generated through the actin cytoskeleton [30]. Munger and colleagues tested a panel of β6-cytoplasmic tail truncations and reported that the β6 mutant that lacked the C-terminal 14 amino-acids was still able to activate latent TGF-β. These data suggest that in our model, both the αvβ6- and αvΔβ6-expressing PDAC cells should be able to activate latent TGF-β at similar levels, since the 11 aa are not required [30]. Thus, a differential activation of latent TGF-β is unlikely to explain the ability of the C-terminal 11aa of to regulate PDAC metastasis to the lung, but additional studies are required.

In summary, we report for the first time that the C-terminal 11 amino acids of the integrin β6 subunit regulates the proliferation, migration and 3D invasion of PDAC cells. Moreover, the same amino-acids regulate the ability of PDAC cells to metastasise to the lung. Since over 90% of patients with PDAC express αvβ6 these results further support that αvβ6 should be considered a major target for anti-metastatic therapy in PDAC.

## Materials and methods

### Cell culture

Murine PDAC tumour cells were derived from KPC: TB32043 (LSL-Kras^G12D/+^; LSL-Trp53^R172H/+^; Pdx-1-Cre) [17] and KPF: TB32048 (FSF-Kras^G12D/+^; LSL-Trp53^frt/+^; Pdx-1-Flp) [18] mice. Cells were cultured in DMEM (41966-029, Gibco) supplemented with 10% Fetal Bovine Serum (FBS) (A5670701, Gibco) and 1% Penicillin/Streptomycin (P/S) (15140-122, Gibco) at 37ºC and 5% (v/v) CO_2_.

Where indicated, cells were treated with αvβ6-blocking antibodies. 10D5 antibody (Sigma-Aldrich) and used in a 1:100 dilution in DMEM 10% FBS. Treatment with 264RAD antibody (a gift from Astra Zeneca) was performed in a final concentration of 20 µg/mL.

### Lentiviral production and transduction of cell lines

Control, β6- and Δβ6-expressing cell lines were generated by lentiviral transduction using the pLV-puro, pLV-puro-mItgb6 and pLV-puro-mΔItgb6 (Vectorbuilder), respectively. Lentiviral particles were produced transfecting the lentiviral plasmids together with pVSV.G (Addgene, 14888) and pPAX2 (Addgene, 35002) into HEK293T cells using Lipofectamine 2000 reagent (11668027, ThermoFisher). After 72h, supernatants were collected and ultracentrifuged (23,000g, 2 h, 4 C). Concentrated lentiviral particles were then added to TB32043 and TB32048 at 70% confluency. Transduced cells were selected by adding puromycin at 1.5 µg/mL final concentration for 72h. β6- and Δβ6-expressing cells were FACS sorted twice (selecting with 10D5 detected by Alexa-488 conjugated anti-mouse IgG-(A-11001, ThermoFisher)) to select the high αvβ6expressing cells.

### Western blotting

For protein analyses, cells were washed two times with cold PBS and cell lysates were prepared using 1% SDS lysis buffer (25mM Tris pH 7.5, 200mM NaCl, 1% SDS). Lysates were incubated for 15 minutes at 95ºC. Protein concentration was measured using the DC™ Protein Assay Kit (5000122, Bio-Rad) following manufacturer’s protocol. 20 µg of protein was loaded into 4–15% Mini-PROTEAN™ TGX Stain-Free™ gel (4568086, Bio-Rad) and the gel was run at 100V for the first 15 minutes and 200V until protein front runs out of the gel. Proteins were transferred to a nitrocellulose membrane (Amersham Hybond^TM^, GE Healthcare) for 1h 30’ at 100V. After transferring, membranes were blocked for 45’ in 1% BSA in TBST at room temperature (RT). Membranes were incubated with primary antibodies overnight at 4ºC. The following antibodies were used: goat anti-N-terminal integrin β6 (detecting both β6 and Δβ6) (AF2389, R&D Systems), goat polyclonal antibody to the integrin β6 C-terminal distal amino-acids (detecting only wild-type β6) (sc6632, Santa Cruz) and β-actin antibody (ab8226, Abcam). Secondary antibodies were incubated for 1h at RT (P0160 and P0260, Dako, Agilent Technologies). Protein detection was performed using the Bio-Rad ChemiDoc imaging system.

### Flow cytometry and FACS sorting

For flow cytometry experiments, cells were detached and dissociated with trypsin/EDTA and washed three times with ice cold flow buffer (DMEM, 0.1% (w/v) BSA and 0.1% (w/v) sodium azide (NaN_3_) (S8032, Sigma Aldrich). Cells were incubated with anti-αvβ6 antibody 10D5 for 1 h at 4ºC. After incubation, cells were washed again with cold flow buffer and incubated with Alexa Fluor 488-conjugated goat anti-mouse secondary antibody (A-11001, ThermoFisher) at a 1:200 dilution for 30 min at 4ºC in the dark. Cells were then washed again in cold flow buffer and resuspended in flow buffer containing 0.5mg/mL DAPI 1:10.000 (62248, ThermoFisher). Samples were analysed by flow cytometry on a BD LSR Fortessa (BD Biosciences).

For FACS sorting, cells were processed in the same way as for flow cytometry experiments. For sorting, cells were finally resuspended in flow buffer with DAPI and 5mM EDTA. After resuspending the cells, these were filtered through a 40µm strainer prior to sorting (CLS431750, Corning). Cells were then FACS sorted twice in BD FACS Aria Cell Sorter (BD Biosciences) to select the highest β6- and Δβ6-expressing cells.

### Proliferation assay

Cell proliferation was measured using the Incucyte live-cell imaging and analysis instrument (Sartorius). 1.000 cells were seeded in 96-well plates in triplicates. After an over-night incubation to allow cells to attach, plates were placed in the Incucyte and imaged for 72h. Proliferation was measured quantifying cell confluency using the Incucyte software. Treatments with αvβ6-blocking antibodies were done prior placing the plates in the Incucyte.

### Migration assay

For cell migration experiments, cells were seeded in 24-well plates in triplicates. A wound was made using a p20 tip when cells were at confluency. Brightfield images of the wounds were taken at time 0h. Cells were incubated for 16h to allow wound closure and images were taken after incubation. Wound area of images at time 0h and 16h was quantified using ImageJ software and represented as percentage of closed wound.

### Spheroid 3D invasion assay

Invasion was measured using the 3D spheroid invasion model described in Coetzee et al 2023 [20]. PDAC cells spheroids were formed in 2.5% (v/v) methylcellulose (M0512, Sigma) hanging drops. For mono-culture spheroids, 1000 cells were used per spheroid. In co-culture experiments, 1000 cells were used in a 2:1 ratio PSC: cancer cell. Spheres were collected the next day and suspended in organotypic mixture (For 1mL of mixture: 2mg/mL Collagen (354249, Corning), 175µL Matrigel, 50µL HEPES (1 M, pH 7.5, H7006, Sigma), 25µL NaOH and DMEM 10% FBS) before seeding into 96-well plate. Culture media was added on top of gels. When indicated, treatments were added also on top of the gels. Spheroids were incubated for 72h and imaged using an Axiovert 135 (Carl Zeiss MicroImaging LLC). Percentage of invasive area was quantified in ImageJ, using the following equation: % invasive area = ((total area − central area)/central area) × 100.

### Quantitative real time PCR (qRTPCR)

RNA was obtained using TRIzol (15596026, ThermoFisher) RNA isolation protocol. RNA was retro-transcribed and analysed by quantitative PCR in triplicates using QuantStudio 7 (Applied Biosystems, Thermo Fisher Scientific). Expression levels were normalised to HPRT expression. PCR primers are listed in Table 1.

**Table 1:**
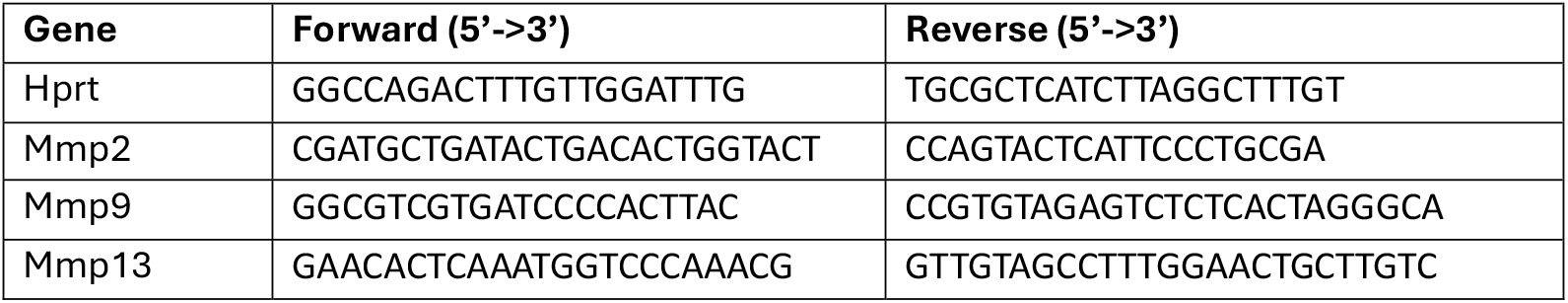
List of primers used for qRT-PCR analyses.

### Statistical analysis

All results presented are representative from at least three independent experiments. Data are shown as mean ± one stabdard deviation (SD). Statistical analysis was performed using GraphPad Prism (Systat Software, USA). For 2 variables, data were analysed using unpaired-two-tailed student t-test. For 3 or more variables data were analysed using one-way ANOVA with Tukey’s multiple comparisons test.

### Orthotopic murine PDAC model

The in vivo experiment was performed in accordance with the guidelines issued by the UK Home Office under approved Project Licenses. C57BL/6J mice (8 weeks old) were used and orthotopic PDAC cells implantation was performed as described in Brown et all 2024 [21]. 1.000 Ctl, β6- and Δβ6-expressing TB32043 or TB32048 cells mixed with Matrigel (354234, Corning) to a volume of 30 µL were injected into the pancreas using an insulin syringe (0.3 ml MicroFine, 324826, BD). The pancreas and spleen were returned to the abdominal cavity, the peritoneum closed with absorbable sutures (C0022002, Braun), and the skin with wound clips (726063, Harvard Apparatus). Wound clips were removed one week after surgery. Tumour growth was measured by IVIS imaging (Ilumina) weekly after day 7 post-injection. Mice were terminated following UK Home Office regulations when the Humane Endpoints (HE) was reached.

## Acknowledgements

GM was funded by a Pancreatic Cancer Research Fund award and DDB was funded by a Medical Research Fund award (Grant Ref: MR/W02537X/1). We are grateful to staff in Barts Cancer Institute Core Services (flow cytometry, microscopy, histology) for their expert assistance and for the Cancer Research UK City of London Centre Grant to Barts Cancer Institute (C355/A25137 XX).

